# Aerobic exercise prevents the loss of endogenous pain modulation in male and female rats with traumatic brain injury

**DOI:** 10.64898/2026.03.31.714901

**Authors:** Karen-Amanda Irvine, Adam R. Ferguson, J. David Clark

**Author notes:** **Corresponding author**: Karen-Amanda Irvine.

## Abstract

Traumatic Brain Injury (TBI) patients may suffer from a number of long-term complications after injury such as impaired motor skills, cognitive decline, and sensory abnormalities including chronic pain. Disruption of endogenous pain modulatory pathways likely contributes to development of chronic pain in a wide range of conditions including TBI. Aerobic exercise has been shown to impact pain syndromes. Here we investigate the effect of exercise on pain outcome measures after TBI using a lateral fluid percussion (LFP) model and voluntary running wheels in male and female rats. We tested mechanical nociceptive reactivity with von Frey fibers and descending control of nociception (DCN) using hindpaw sensitization with PGE2 followed by a capsaicin-test stimulus to the forepaw. Pharmacological studies employed the administration of noradrenergic (NA) and serotoninergic receptor blockers. Neuropathological studies quantified neuroinflammatory changes and axonal damage. We found that exercise decreased the duration of the acute phase of pain from ∼5 weeks to 2-3 weeks in female and male TBI rats respectively, gains that could be reversed using the α_1_-adrenoceptor (α_1_AR) antagonist, prazosin. Exercise also prevented the loss of DCN for at least 180 days post-injury in both male and female TBI rats. The intact DCN response in male and female TBI rats provided by exercise could be blocked using prazosin. Surprisingly, exercise-mediated restoration of the DCN response in male TBI rats was not blocked by the 5-HT_7_ receptor antagonist, SB-267790, the receptor system through which serotonin reuptake inhibitors restore DCN after TBI in male rats. Therefore, the transition from a noradrenergic to a serotonergic inhibitory pain pathway that we typically see in male TBI rats, was blocked by exercise. Assessment of neuropathology, acutely after TBI, reveals that both the astrocyte and microglial response to injury is significantly greater in male TBI compared to female TBI, regardless of exercise. The effect of exercise on the extent of neuroinflammation after injury was minimal in TBI rats of both sexes. In contrast, exercise significantly decreased the amount of axonal loss in the corpus callosum in both male and female TBI rats compared to sedentary TBI rats. However, the extent of axonal loss after TBI in both exercise and sedentary male rats was greater than in female exercise and sedentary groups respectively. These results demonstrate that exercise is a promising treatment for chronic pain after TBI in both male and females. It also highlights that dysfunction of the endogenous pain modulatory pathways observed in male rats after TBI can be prevented by exercise, possibly by reducing axonal loss.

## Introduction

Traumatic brain injury (TBI) is a major public health issue impacting approximately 60 million individuals each year globally^1^. Defined as an injury to the brain by an external force, TBI remains the leading cause of death and disability among all trauma-related injuries worldwide^2,3^. TBI patients frequently experience sensory issues, impaired motor control, emotional instability and chronic pain^4–6^. Chronic pain in the setting of TBI contributes to disability, causes suffering, complicates rehabilitative efforts and poses a major challenge to clinical management teams^7^. The most frequently endorsed types of pain include headache, back and musculoskeletal/joint pain. As of yet, specific treatments for TBI-related pain remain elusive.

While there are a number of potential candidates that could be responsible for the pain perceived post-TBI (i.e. endocrine system abnormalities or sleep disturbances), dysregulation of descending endogenous pain modulatory pathways from the brain to spinal cord is one of the best-supported explanations ^8^. Experimental evidence from our team and others suggests that TBI causes damage to endogenous pain control centers in the brainstem resulting in diminution of descending pain inhibition and/or enhancement of descending pain facilitation. This then supports the development of central sensitization in the spinal cord, a key feature present in other chronic pain disorders ^9,10^.

Our work using rodent models of TBI demonstrate failure of descending pain modulation, referred to as descending control of nociception (DCN) when measured in conscious behaving animals^11^. The neurophysiological basis of DCN failure following TBI in rats revealed changes in nociceptive signaling after injury manifests in two distinct phases. The initial stage is characterized by hindlimb hyperalgesia that occurs within 24 hours of injury and resolves after 4-5 weeks post-injury. For both males and females, restoration from this stage is dependent on either a reduction of descending serotonergic pain facilitation via 5-HT_3_ receptor antagonism or via the enhancement of descending noradrenergic (NA) pain inhibition by increasing the spinal levels of NA with the NA reuptake inhibitor, reboxetine (RBX)^12,13^. The adrenoceptor responsible for the descending NA pain inhibitory pathway in naïve rats and rats after peripheral nerve injury has been consistently shown to be the α_2-_adrenoceptor (α_2-_AR) ^14,15^. However, we have shown that after TBI the descending NA inhibitory pathway switches from an α_2_-AR to α_1_-AR driven mechanism by 7 DPI^13^. Also present at this acute stage are significant neuroinflammatory changes that occur within several key areas of the brain and spinal cord which are involved in descending pain modulation^16^. Resolution of the acute stage of pain after TBI then gives rise to the chronic stage, more relevant to persistent pain, that is characterized by a failure of the DCN response and which is sustained for up to at least 24 weeks post-injury^16^. In females, the DCN response can be restored pharmacologically by enhancing descending NA inhibitory signaling via RBX through the α_1_-AR^13^. In contrast, restoration of DCN response in males requires enhancement of descending serotonergic inhibition using the selective serotonin reuptake inhibitor, escitalopram (ESC) via the 5-HT_7_ receptor ^13,17^ ^18^.

A promising candidate therapy for the amelioration of post-TBI pain is aerobic exercise. In fact, exercise is a treatment recommendation central to chronic pain management, particularly in the management of musculoskeletal conditions such as joint and back pain as well as widespread pain conditions such as fibromyalgia ^19^. Our group has shown that voluntary exercise can limit early sensitization and prevent the loss of DCN after a closed head model of TBI that mimics a mild concussive injury. In addition, we have shown that exercise limits the spinal cord expression of pronociceptive genes after TBI and augments injury-induced neuroinflammation^20^. Here, we advance our studies on chronic pain after TBI, by investigating the role of voluntary wheel running exercise in Sprague-Dawley male and female rats after a lateral fluid percussion injury and identify the neurotransmitter receptor mechanism responsible for those effects.

## Materials and Methods

Greater detail on the materials and methods used in these studies may be found in our previous publications. ^12,14,16,18^

### Animals

All studies were approved by the Veterans Affairs Palo Alto Health Care System Institutional Animal Care and Use Committee (Palo Alto, CA, USA) and followed the animal guidelines of the National Institutes of Health Guide for the Care and Use of Laboratory animals (NIH Publications, 8^th^ edition, 2011)^21^. Male (n=72 250-300g) and female (n=72 225-275g) Sprague Dawley rats (Inotiv, Indianapolis, IN, USA), were housed under standard conditions with a 12h light–dark cycle (6:30 am to 6:30 pm) and were given food and water *ad libitum*. The animals were housed in pairs in 30 × 30 × 19-cm isolator cages with solid floors covered with a 4 cm layer of wood chip bedding. Experimenters were blinded to the identity of treatments and experimental conditions. All studies were designed to minimize the number of rats required. All *in vivo* experiments were performed between 6 am and 2 pm in our facility’s Veterinary Medical Unit.

### Drugs

Prostaglandin E2 (PGE-2) [1μg/50μl], intraplantar (*i.pl.*) (14010, Cayman Chemicals, MI, USA) a principal mediator of inflammation and pain hypersensitivity. Capsaicin (CAP) [10μg/10μl], *i.pl.* (M2028, Millipore-Sigma, MO, USA) that causes hyperalgesia to mechanical stimuli at and around the injection site. Reboxetine Mesylate (RBX) [40mg/kg], *i.p.,* (R1126, Ontario Chemicals, Canada) a noradrenergic reuptake inhibitor (NRI). Escitalopram hydrochloride (ESC) [10 mg/kg] *i.p.,* (462.340.010, Acros Organics, PA USA) a selective serotonin reuptake inhibitor (SSRI). Atipamezole hydrochloride (ATZ) [1 mg/kg], *s.c*., (9001181, Cayman Chemicals) an alpha-2-adrenoceptor antagonist. Prazosin hydrochloride (PRZ) [1mg/kg], *i.p,* (15023, Cayman Chemicals, MI, USA) an alpha-1-adrenoceptor antagonist. SB-269970 (SB-269970) [10µg/10µl], i.t., (17081, Cayman Chemicals) a 5-HT_7_ receptor antagonist. All drugs were diluted in sterile saline and prepared on the day of testing. The doses of these drugs were chosen based on previous studies ^22^.

### Running Wheel Exercise Protocol

Rats had ad libitum access to the running wheels (Starr Life Sciences Corp, PA, USA) in their cages for 24h/day, 7 days a week for 2 weeks. On the day of injury, the running wheels (35 cm in diameter) were locked and nonfunctional for a period of 3 days (**Figure 1A**). This was based on previous work revealing negative effects of initiating exercise too soon after rodent TBI^20,23^. On day 3 after TBI, rats were randomly assigned to either the exercise group which had their cage wheels unlocked or the sedentary group in which the cage wheels remained locked for the remainder of the exercise wheel protocol of up to 7 weeks post-TBI. Running wheels were connected to a manual LCD counter which was checked and recorded daily to monitor the number of wheel revolutions achieved that day. To obtain the distance in meters (m), the diameter of the wheel in meters (0.35 m) was multiplied by π to obtain the circumference and this value was multiplied by the number of revolutions (**Figure 1B**). On day 49, after DCN testing, all rats were removed from exercise cages and singly housed in standard rat filter cages until 180 DPI.

**FIGURE 1:**
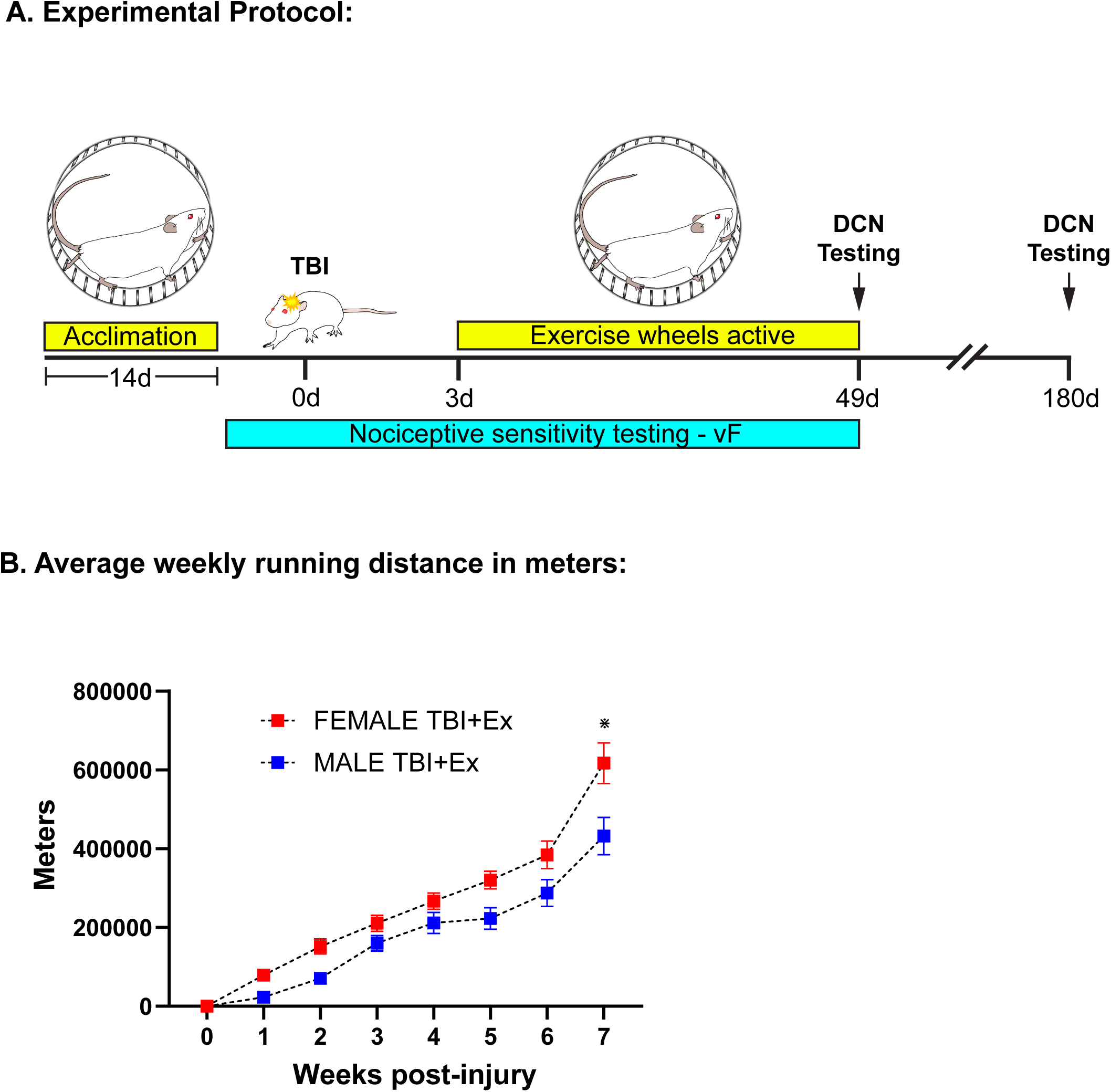
A schematic diagram of the running wheel exercise protocol used in this study. All rats were acclimated to functional running wheels 24 hours a day, for a total of 14 days (**Fig 1A**). On day 15, all rats received a lateral fluid percussion injury to the right side of the head. Following surgery, the injured rats are returned to the running wheels cages with the wheels locked for a total of 3 days. After 3 days, the wheels for half the rats were unlocked to create the TBI and exercise group (TBI+Ex) and the wheels for the remaining rats remained locked to create the TBI and sedentary group (TBI+Sed). Von Frey analysis was conducted prior to the TBI and regularly throughout the study up until 49 days post injury (**Fig 1A**). The descending control of nociception response was assessed using the noxious stimulation–induced anti-nociceptive protocol on days 49 and 180 after injury. Note: rats were removed from running wheel cages and returned to standard filtered rat cages after DCN testing on day 49 until sacrifice. **Fig 1B**: The average weekly distance travelled by female and male TBI+Ex rats. No significant difference was found up to 6 weeks post-injury, however at 7 weeks post-injury, female TBI+Ex rats ran significantly more than male TBI+Ex rats (*p < 0.01). Data are presented as mean + standard error of the mean (n = 6 rats per cohort).

### TBI surgery

Rats were anesthetized with isoflurane throughout the procedure (5% for induction and 2% for maintenance) and the paw pinch reflex was used to ensure the state of anesthesia. The lateral fluid percussion (LFP) model of TBI was used as described previously ^12,16,18^. Briefly, a midline incision was made in the scalp, and a 5 mm craniotomy was made on the right side of the skull using a mini-drill and a 5 mm trephine burr (Fine Science Tools). The craniotomy was placed midway between the bregma and lambda sutures and, 2 mm right of the midline suture. Using cyanoacrylate glue and dental acrylic, a female luer attachment was affixed to the craniotomy opening. Following recovery of pinch reflexes, the luer attachment was connected to the LFP apparatus (Amscien Instruments, VA, USA), and a pressure wave of 1.3 ±0.1 atm to produce mild level injuries or no pressure wave (sham) was applied to rat dura based on previous reports. Thereafter, the luer attachment and dental acrylic were removed, the bone flap replaced, and the overlying wound closed using staples. Rats were allowed to recover on a warming pad prior to being returned to their home cage

### Intrathecal injection

Rats were anesthetized with isoflurane procedure (5% for induction and 2% for maintenance) and a 3 cm^2^ window of fur, near the base of the tail, was shaved and washed with 70% ethanol to maximize visualization during needle insertion. An empty 50ml conical centrifuge tube was placed underneath the rat to arch the lumbar vertebral column, and the spinous process of L5 was located. The spinous process of L5 was pulled in a cranial direction and the vertebral body of L6 was pulled caudally towards the tail. This maximized the space within the groove between L5 and L6 vertebrae into which a 25G needle was carefully inserted. Successful entry of the needle into the intradural space was confirmed by the observation of a tail flick. The needle was immediately secured into position with one hand and the desired volume of substance slowly injected with the other hand. Once the injection was performed, the rat was placed on a warming pad prior to returning to their home cage.

### Behavioral testing

All behavioral assessments were performed by observers who were blinded to the experimental conditions. Mechanical withdrawal thresholds were measured using a modification of the up-down method and von Frey filaments as described previously^24^. Animals were placed in clear Plexiglas boxes (17cm length x 11cm width x 13cm height) on an elevated horizontal wire mesh stand (IITC Life Science Inc, CA, USA). After 60 minutes of acclimation, fibers of increasing stiffness with initial bending force of 4.31N (ranging from 4.31N up to 5.18N) were applied to the plantar surface of the hindpaw and left in place for 5 seconds with enough force to slightly bend the fiber. Withdrawal of the hindpaw from the fiber was scored as a response. When no response was obtained, the next firmer fiber in the series was used in the same manner. If a response was observed, the next less stiff fiber was applied. Testing continued until 4 fibers had been applied after the first one causing a withdrawal response, allowing the estimation of the mechanical withdrawal threshold using a curve fitting algorithm ^25^.

### DCN paradigm

Assessment of post traumatic mechanical hypersensitivity was done using a variant of a widely-used noxious stimulation–induced anti-nociceptive protocol ^26–28^. Briefly, the rats were confirmed to recover to baseline mechanical thresholds after TBI (at 49 days post-injury). Next, an intraplantar *(i.pl.*) injection of PGE-2 (conditioning stimulus) into the hindpaw contralateral to the TBI was used to produce brief hypersensitivity ^29^. An hour after the PGE-2 injection, hindlimb withdrawal thresholds were measured using the von Frey filaments as described above. Subsequent to mechanical threshold evaluations, rats were given a capsaicin injection (test stimulus) into the dorsal surface of the forepaw ipsilateral to the TBI to induce DCN. All *i.pl* injections were administered under light isoflurane anesthesia. At 15, 30, 60, 120, 180 and 240 min after capsaicin injection, withdrawal thresholds were measured using the von Frey filaments as described above.

### Immunohistochemistry

This protocol has been described in detail previously ^12,16^. Briefly, rats were perfused with ice cold PBS and then 4% paraformaldehyde, brains were then removed and cryoprotected in 20% sucrose in phosphate buffered saline (PBS) for 2 days at 4°C. The brains were cut into 15 µm transverse sections using a cryostat. Immunostaining was performed using antibodies against rabbit anti-glial fibrillary acidic protein for astrocytes (GFAP; 1:1000, AB5804, Millipore-Sigma), rabbit anti-ionized calcium binding adaptor molecule-1 for microglia (IBA-1, 1:750, 019–19,741, Wako, VA, USA), and mouse monoclonal antibody to non-phosphorylated neurofilaments for axons (SMI-31R, 1:1000, 801601, Biolegend). Blocking of all the sections took place at room temperature for 2 hours in PBS containing 10% normal goat serum (Vector Laboratories, CA, USA), followed by exposure to the primary antibody overnight at 4°C. Controls prepared with the primary antibody omitted showed minimal background fluorescence under the conditions employed. Staining was performed concurrently for each group of sections compared with one another and photographed under identical conditions.

### Image analysis

In all data assessments, both the photography and image analysis were performed by observers who were blinded to the experimental conditions. Expression of GFAP and IBA-1 was evaluated bilaterally in the dentate gyrus (A–P −2.0 to −4.5mm), thalamus (A–P −3.0 to-5.0mm), somatosensory cortex 1, callosal radiations (CR) (A-P −2.0 to −4.5mm), periaqueductal gray matter (PAG, A–P −6.0 to −8.75 mm), RVM (A-P −10.8 to −12.8mm), and locus coeruleus (LC, A-P −9.0 to −10.5mm) on four coronal sections per brain spaced 150μm apart using a Keyence All-in-One Fluorescence Microscope (BZ-X700, Keyence Corporation of America, Itasca, IL) with appropriate filter sets. Sections from uninjured, untreated rats were used to establish a threshold level that excluded all nonspecific staining. This threshold level was then applied to all experimental groups. The percentage of the total image area covered by GFAP and IBA-1 in each section was calculated using the ‘‘area fraction’’ feature in the NIH ImageJ program Axonal loss was quantified in the medial corpus callosum in four coronal sections per brain, spaced 150 μm apart, immunostained for SMI31R. The relative number of axons in the corpus callosum was estimated using a point grid overlaid on the stained sections taken at x40 magnification. The axon density was calculated by counting the number of grid points that fell on an axon, subtracting this value from the total number of grid points, and multiplying by 100 to express this as a percentage. This percentage was then multiplied by the dorsal-medial thickness of the corpus callosum to derive the relative number or axons^16,30^.

### Data analysis

An independent statistician (A.R.F.) tested statistical assumptions, performed missing values analysis, descriptive statistical tests, followed by inferential tests. Data met normality, homogeneity of variance, and independence assumptions sufficiently for application of general linear models (GLM). Group comparisons were therefore performed according to *a priori* balanced experimental designs as 2- 3-, or 4-way, mixed repeated measures ANOVAs, modeling between- and within-subject factors as appropriate using GLM. Significant effects were followed by interaction plots and Bonferroni’s post-hocs. All statistical analyses were performed using IBM SPSS software (v.31, IBM Corp.) with base, regression, missing values, and advanced statistics modules. Statistical code is freely available upon request. Graphs were created using Prism 10.6.0 (GraphPad Software). Data are presented as mean values +/- standard error of the mean (SEM). All experimental sample sizes (n) were selected by *a priori* power calculations based on historical data from our laboratory. The figure legends report precise F values, degrees of freedom, P values, effect sizes, and observed power for all significant effects as well as the n’s completed for each group.

### Data availability

All the raw data are available through the VA- and NIH-supported open data commons for TBI (odc-tbi.org), in support of funder mandates for FAIR (findable, accessible, interoperable, and reusable) data stewardship policies. Data may be accessed publicly through the Open Data Commons for Traumatic Brain Injury (odc-tbi.org) at the following persistent digital object identifier: doi.org/10.34945/F5F010. The dataset can be reused with attribution to the original data creators via the creative commons 4.0 BY license.

## Results

### Running wheel performance after TBI is similar between male and female rats

The average distance travelled in meters was compared between male and female TBI plus exercise (TBI+Ex) rats every week after injury for 7 weeks. A two-way mixed repeated measures ANOVA revealed that there was no significant difference in the distances ran between male and female TBI+Ex rats for up to six weeks post-injury (**Figure 1B**). However, *post hoc* analysis revealed that at 7 weeks post-injury female TBI+Ex rats ran significantly more than the male TBI+Ex rats (**Figure 1B**) (*p < 0.01).

### Exercise significantly reduces the duration of the acute pain phase in both male and female TBI rats

All male and female TBI rats, regardless of exercise, displayed injury-induced mechanical allodynia in the contralateral and ipsilateral hindpaws that reached maximal severity by 3 days post-injury (DPI) (**Figure 2A** and **2B**). At 7 DPI, female exercised-TBI (TBI+Ex) rats displayed a significant decrease in both ipsilateral and contralateral hindpaw hypersensitivity compared to female sedentary-TBI (TBI+Sed) rats (**Figure 2A**). This is in contrast to male TBI+Ex rats which showed no change in hypersensitivity of either hindpaw at 7 DPI compared to male TBI+Sed rats (**Figure 2B**). By 14 DPI, hindpaw withdrawal threshold had returned to pre-injury levels in female TBI+Ex rats. In contrast, although hindpaw hypersensitivity had significantly decreased compared to male TBI+Sed rats, male TBI+Ex rats did not achieve preinjury hindpaw sensitivity levels until day 21 post-injury (**Figure 2A** and **2B**). The contralateral and ipsilateral hindpaws of all the TBI+Sed rats, regardless of sex, remained hypersensitive up until day 35 post-injury. A four-way mixed repeated measures ANOVA revealed significant main effects of exercise (p=2.08 x 10^-14^), sex (p=0.028), laterality (ipsilateral versus contralateral) (p=6.50 x 10^-3^) and time post-TBI (p=2.23 x 10^-98^). There were also significant interactions of exercise by sex (p=1.57 x 10^-7^), laterality by sex (p=1.49 x 10^-4^), laterality by exercise (p=0.0197), time post-TBI by sex (p=1.08 x 10^-10^), time post-TBI by exercise (p=5.65 x 10^-38^), time post-TBI by laterality (p=2.04 x 10^-15^), laterality by exercise by sex (p=3.83 x 10^-5^), sex by time post-TBI by exercise (p=1.89 x 10^-22^), sex by time post-TBI by laterality (p=4.20 x 10^-18^), exercise by time post-TBI by laterality (p=3.47 x 10^-12^) and sex by exercise by time post-TBI by laterality (p=7.02 x 10^-18^). *Post hoc* analysis reveals that the male and female TBI+Ex groups had significantly higher pain thresholds than the TBI+SED groups of the same sex (*p < 0.001).

**FIGURE 2:**
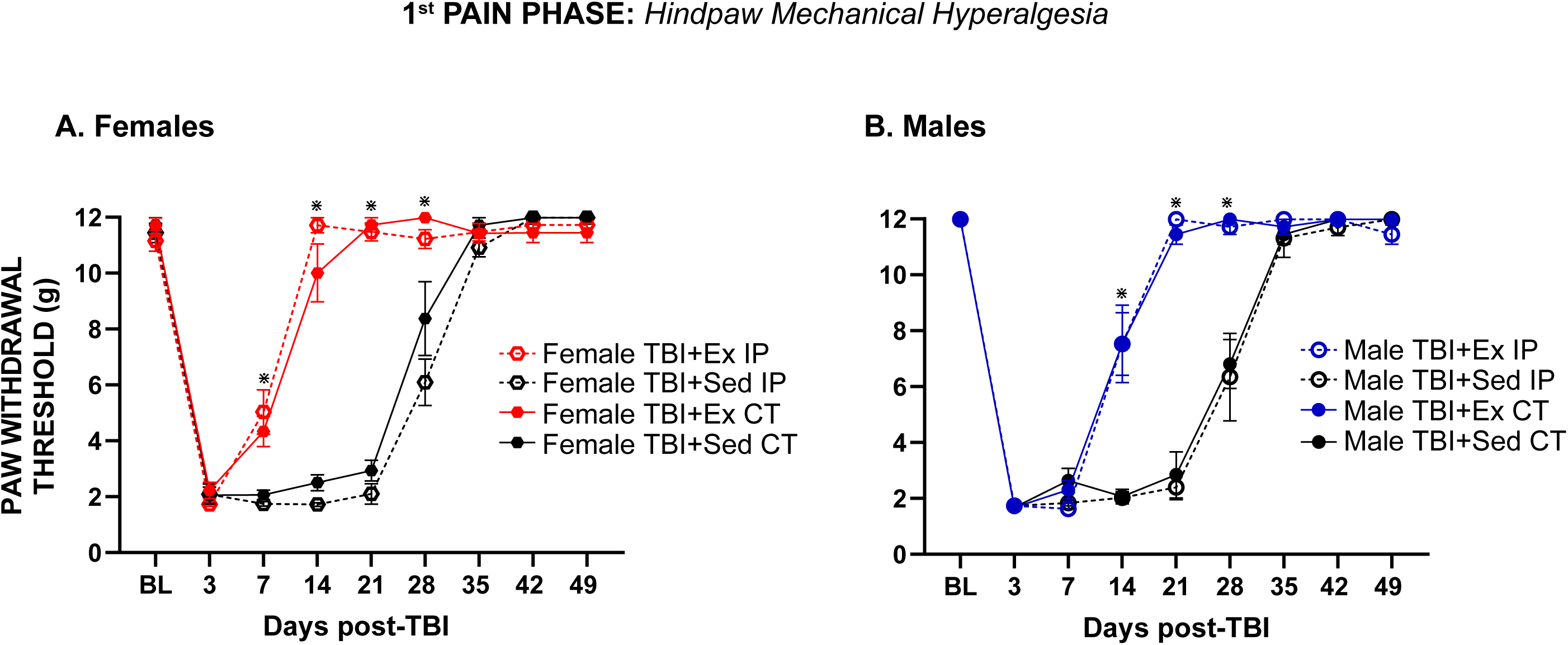
The effect of exercise on the acute stage of pain in both female (**Fig 2A**) and male (**Fig 2B**) TBI+Ex and TBI+Sed rats. The acute stage of pain is characterized by mechanical hypersensitivity of both the contralateral (CT) and ipsilateral (IP) hindlimbs within 24 hours of injury. Females showed a significant increase in paw withdrawal threshold (PWT) of both the CT and IP hindlimbs at 7 days post-injury (DPI) when compared to female TBI+Sed rats (**Fig 2A**). Preinjury hindlimb sensitivity in female TBI+Ex rats returns by 14 DPI when compared to female TBI+Sed rats (**Fig 2A**). Also, at this time there is a significant increase in PWT of hindlimbs in the male TBI+Ex rats compared to the TBI+Sed rats (**Fig 2B**). Normal PWT is reestablished in the male TBI+Ex rats by 21 DPI (**Fig 2B**). Mechanical hypersensitivity persisted up until 28 DPI in both male and female TBI+Sed rats and returned to preinjury levels by 35 DPI. (**Fig 2A** and **2B**). A four-way mixed repeated measures ANOVA revealed significant main effects of exercise (F(1,20)=373.41, p=2.08 x 10^-14^, □2=0.95), sex (F(1,20)=5.64, p=0.028, □2=0.22), laterality (ipsilateral versus contralateral) (F(1,20)=9.21, p=6.50 x 10^-3^, □2=0.32) and time post-TBI (F(7,140)=547.96, p=2.23 x 10^-98^, □2 =0.97). There were also significant interactions of exercise by sex (F(1,20)=61.58, p=1.57 x 10^-7^, □2=0.76), laterality by sex (F(1,20)=21.77, p=1.49 x 10^-4^, □2=0.52), laterality by exercise (F(1,20)=6.42, p=0.0197, □2=0.24), time post-TBI by sex (F(7,140)=10.64, p=1.08 x 10^-10^, □2=0.35), time post-TBI by exercise (F(7,140)=57.16, p=5.65 x 10^-38^, □2= 0.74), time post-TBI by laterality (F(7,140)=16.10, p=2.04 x 10^-15^, □2= 0.45), laterality by exercise by sex (F(1,20)=27.62, p=3.83 x 10^-5^, □2=0.58), sex by time post-TBI by exercise (F(7,140)=25.86, p=1.89 x 10^-22^, □2=0.56), sex by time post-TBI by laterality (F(7,140)=19.57, p=4.20 x 10^-18^, □2=0.50), exercise by time post-TBI by laterality (F(7,140)=12.28, p=3.47 x 10^-12^, □2=0.38), and sex by exercise by time post-TBI by laterality (F(7,140)=19.27, p=7.02 x 10^-18^, □2=0.49). ⋇ = TBI+Ex hindpaw vs. TBI+Sed hindpaw (p < 0.001), by Bonferroni’s posthocs. Data are presented as mean ± standard error of the mean (n = 6 rats per cohort).

### Exercise-mediated restoration of the acute phase is dependent on intact α1 adrenoceptor signaling in both male and female TBI rats

As described previously, the descending NA inhibitory pain pathway switches from a α_2_-adrenoceptor (AR) to a α_1_-AR dependent mechanism acutely after TBI (7 DPI)^13^. Our next objective was to see which adrenoceptor was responsible for the exercise-mediated restoration of the acute pain phase. At 21 DPI, by which time the hindlimb sensitivity of both male and female TBI+Ex rats have returned to preinjury levels (**Figure 3A** and **3B**), they were treated with the α_1_-AR antagonist, prazosin. Prazosin (PRZ) reversed the exercise-induced restoration of the acute pain in both male and female TBI+Ex rats when compared to vehicle-treated, male and female TBI+Ex rats (**Figure 3A** and **3B**). A four-way mixed repeated measures ANOVA revealed a significant main effect of drug condition (p=1.10 x 10^-26^). No other main effects or interactions reached significance. *Post hoc* analysis reveals that the male and female TBI+Ex groups treated with PRZ had significantly lower pain thresholds than vehicle-treated TBI+Ex groups of the same sex (*p < 0.001).

**FIGURE 3:**
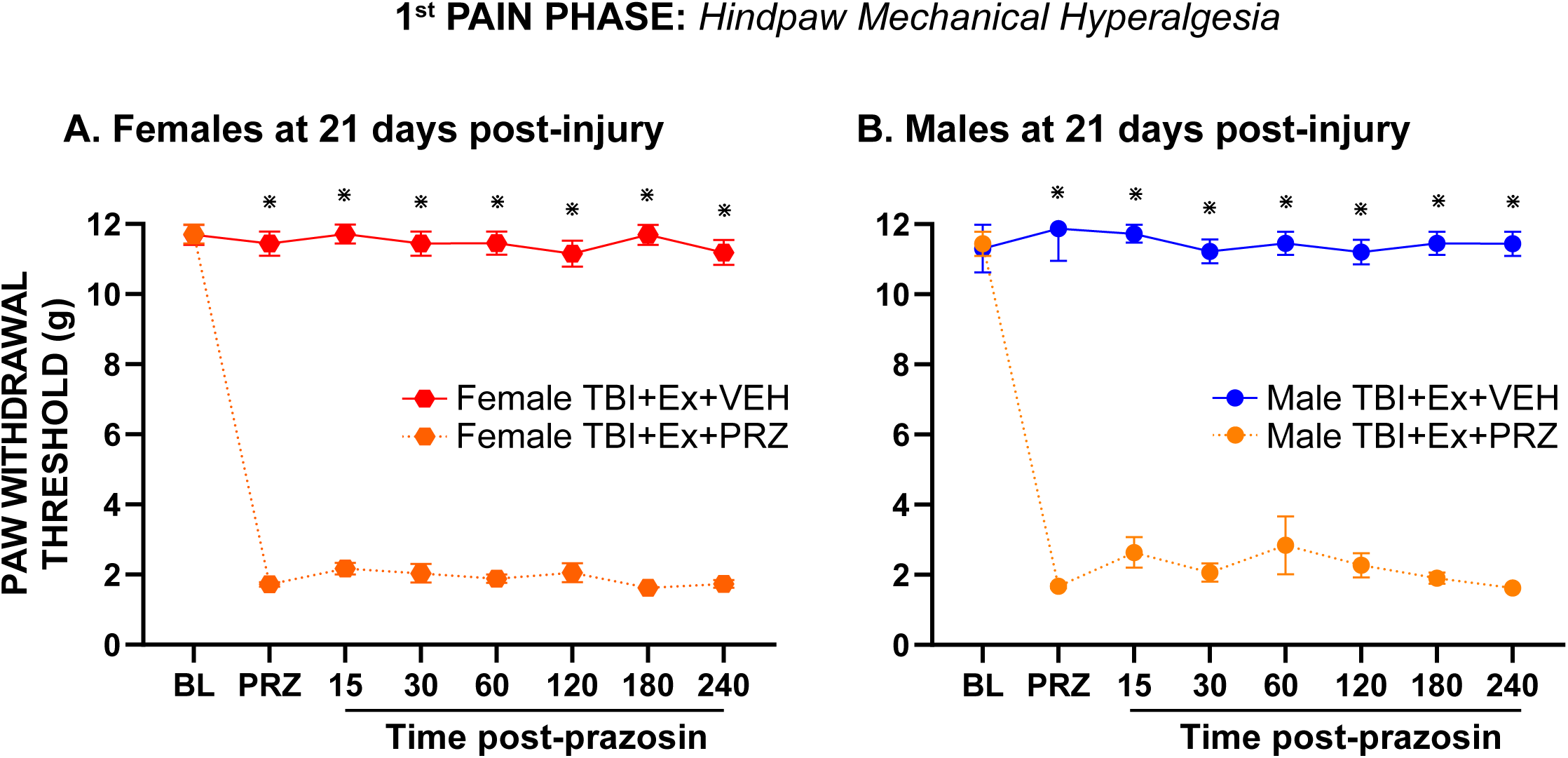
The effect of the α_1_-adrenoceptor (α_1_-AR) antagonist, prazosin (PRZ) on mechanical hyperalgesia in female TBI+Ex rats (**Fig 3A**) and male TBI+Ex rats (**Fig 3B**) at 21 days post-injury (DPI). The exercise-induced restoration of normal hindpaw sensitivity in both male and female TBI+Ex rats by 21 DPI was blocked by treatment with PRZ when compared to VEH-treated TBI+Ex male and female rats. A four-way mixed repeated measures ANOVA revealed a significant main effect of drug condition (F(1,20)=6610.40, p=1.10 x 10^-26^, □2=0.997). ⋇ = TBI+Ex+PRZ vs. TBI+Ex+VEH rats (p < 0.001), by Bonferroni’s posthocs. Data are presented as mean ± standard error of the mean (n = 6 rats per cohort).

### Spinal α_2_-adrenoceptors are critical for descending control of nociception response in both male and female naïve Sprague Dawley (SD) rats

The descending NA pain inhibitory pathway has been consistently shown to be mediated via the α_2_-adrenoceptors (α_2_-AR) in naïve and peripherally injured rats^15,18,31^. We have previously demonstrated that treatment of naïve male SD rats with atipamezole, an α_2_-AR antagonist, will block the DCN response^18^. This verifies the importance of spinal α2AR for noradrenergic-mediated descending pain inhibition when using our noxious stimulation–induced anti-nociceptive protocol in male naïve SD rats^18^. We were able to replicate those experiments (**Figure 4**) and confirm that the DCN response is blocked in both male and female naïve rats treated with atipamezole when compared to vehicle-treated naïve male and female rats (**Figure 4**). A three-way mixed repeated measures ANOVA revealed significant main effects of drug condition (p=3.63 x 10^-4^), and time post-capsaicin (p=6.51 x 10^-27^). There were also significant interactions of drug condition by time post- capsaicin (p=3.66 x 10^-17^), and sex by drug condition by time post-capsaicin (p=3.74 x 10^-4^). *Post hoc* analysis reveals that the male and female naïve groups treated with ATZ had significantly lower pain thresholds than the vehicle treated naïve groups of the same sex (*p < 0.001).

**FIGURE 4:**
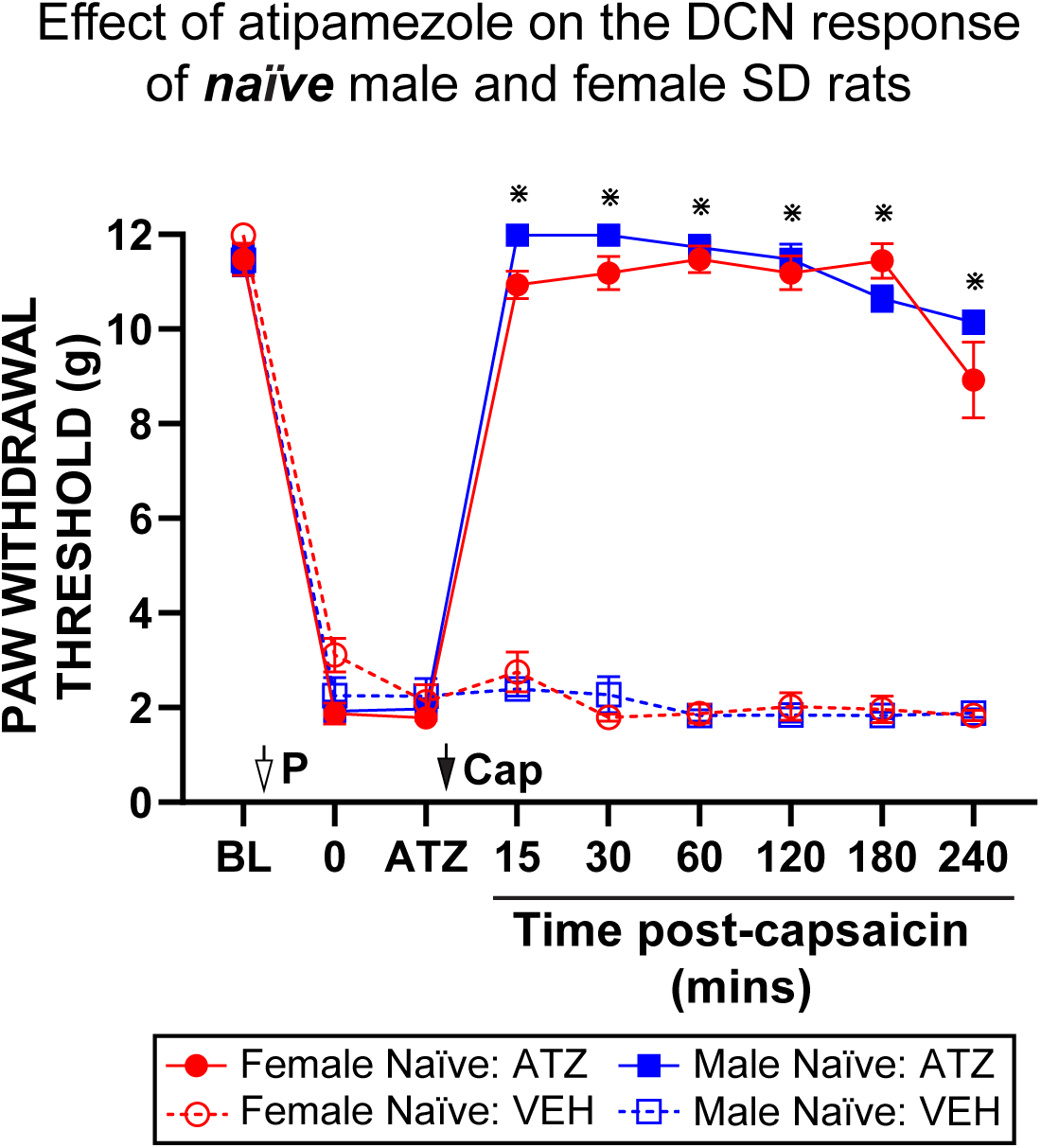
The effect of the α_2_-adrenoceptor (α_2_-AR) antagonist, atipamezole (ATZ) on the intact DCN response of naïve male and female Sprague Dawley rats. The DCN response was blocked in both male and female naïve rats after treatment with ATZ when compared to VEH-treated naïve rats. A three-way mixed repeated measures ANOVA revealed significant main effects of drug condition (F(1,20)=18.34, p=3.63 x 10^-4^, □2=0.48), and time post-capsaicin (F(7,140)=33.31, p=6.51 x 10^-27^, □2=0.63). There were also significant interactions of drug condition by time post-capsaicin (F(7,140)=18.32, p=3.66 x 10^-17^, □2=0.48), and sex by drug condition by time post-capsaicin (F(7,140)=4.13, p=3.74 x 10^-4^, □2=0.17). Open arrow - PGE-2; i.pl. [1 μg/50 μl]. Closed arrow - Capsaicin; i.pl. [10 μg/10 μl]. Data are presented as mean ± standard error of the mean. (n = 6 rats per cohort).

### The descending control of nociception response is intact in both male and female exercised-TBI rats at 49- and 180-days post-injury

To study endogenous nociceptive control systems in the chronic timeframe after TBI, DCN assessment was performed using a noxious stimulation–induced analgesia protocol^27^. The data showed that TBI induces a profound loss of DCN in both male and female TBI+Sed rats at 49 and 180 DPI (**Figure 5A** and **5B** and **Fig. 5C** and **5D**). Conversely, the DCN response in both male and female TBI+Ex rats was intact at both time points (**Figure 5A** and **5B** and **Figure 5C** and **5D**). A four-way mixed repeated measures ANOVA revealed significant main effects of exercise (p=3.13 x 10^-23^), time post-capsaicin (p=2.20 x 10^-48^) and time point (49 or 180 DPI) (p=0.0343). There was also a significant interaction of time post-capsaicin by exercise (p=3.77 x 10^-48^). *Post hoc* analysis reveals that the male and female TBI+Ex groups had significantly higher pain thresholds than the TBI+SED groups of the same sex (*p < 0.001).

**FIGURE 5:** Resolution of the acute phase of pain after TBI exposed the chronic phase of pain involving the failure of descending control of nociception (DCN) from 49 DPI (**Fig 5A** and **5C**) and up to 180 DPI (**Fig 5B** and **5D**) in both male and female TBI+Sed rats. In contrast the DCN response was intact in both male and female TBI+Ex rats at both 49 and 180 DPI (Fig 5). A four-way mixed repeated measures ANOVA revealed significant main effects of exercise (F(1,20)=2974.00, p=3.13 x 10^-23^, □2=0.99), time post-capsaicin (F(6,120)=120.52, p=2.20 x 10^-48^, □2=0.86) and time point (49 or 180 DPI) (F(1,20)=5.16, p=0.0343, □2=0.21). There was also a significant interaction of time post-capsaicin by exercise (F(6,120)=119.26, p=3.77 x 10^-48^, □2=0.86). ⋇ = TBI+Ex CT hindpaw vs. TBI+Sed CT hindpaw (p < 0.0001) by Bonferroni’s posthocs. Open arrow - PGE-2; i.pl. [1 μg/50 μl]. Closed arrow - Capsaicin; i.pl. [10 μg/10 μl]. Data are presented as mean ± standard error of the mean. (n = 6 rats per cohort).

### The presence of a DCN response in female exercised-TBI rats is dependent on intact α1 adrenoceptor signaling

Previous experiments demonstrated that restoration of the DCN response in female TBI rats can be achieved through augmentation of descending noradrenergic antinociceptive signaling in the spinal cord by pharmacologically reducing noradrenalin reuptake^13^. Furthermore, we found that this was an α1 adrenoceptor (α1AR) dependent mechanism^13^. To determine whether exercise works through a similar mechanism, female TBI+Ex rats were treated with prazosin. Prazosin was administered an hour after PGE-2 (conditioning stimulus) and just prior to the test stimulus (capsaicin). Prazosin treatment had no effect on hindpaw sensitivity 15 minutes after its injection *prior* to the test stimulus (**Fig. 6A: PRZ** and **6B: PRZ**). However, following the application of the test stimulus, prazosin blocked the DCN response in the female TBI-Ex at both 49 DPI and 180 DPI (**Fig. 6A** and **6B**). At 49 DPI a four-way mixed repeated measures ANOVA revealed significant main effects of drug condition (p=1.63 x 10^-11^), and time post-capsaicin (p=5.55 x 10^-35^). There was also a significant interaction of time post-capsaicin by drug condition (p=1.13 x 10^-34^). At 180 DPI a four-way mixed repeated measures ANOVA revealed significant main effects of drug condition (p=3.60 x 10^-12^), and time post-capsaicin (p=2.31 x 10^-38^). There was also a significant interaction of time post-capsaicin by drug condition (p=4.74 x 10^-38^). *Post hoc* analysis reveals that the male and female TBI+EX+prazosin groups had significantly lower pain thresholds than the TBI+EX+Vehicle groups of the same sex (*p < 0.001).

**FIGURE 6:**
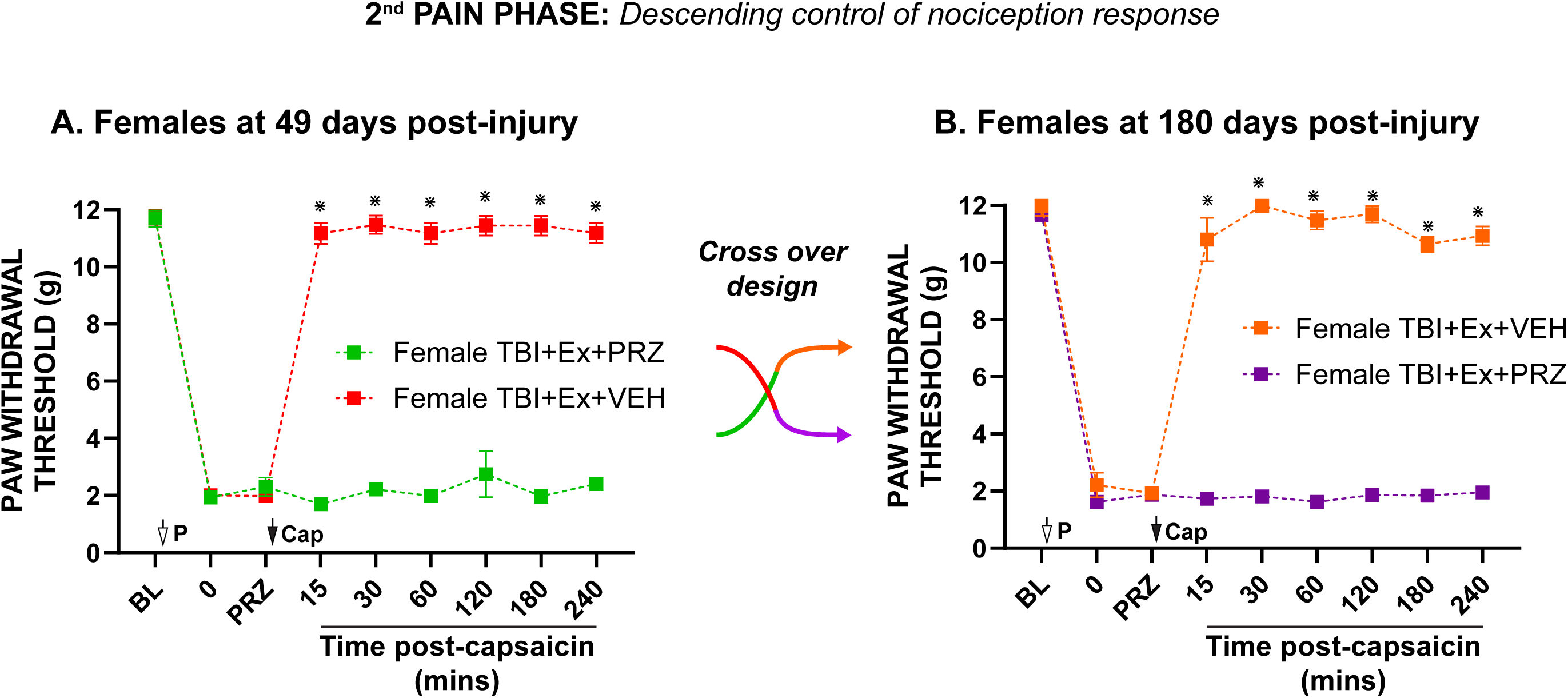
The effect of the α_1_-adrenoceptor (α_1_-AR) antagonist, prazosin (PRZ) on the intact DCN response in female TBI+Ex rats 49 (**Fig 6A**) and 180 DPI (**Fig 6B**). On day 49 of testing, half female TBI+Ex rats were randomly selected and given a systemic injection of PRZ or saline, the vehicle (VEH) (**Fig 6A** and **6B**) after the test stimulus (PGE-2). The DCN response was blocked in all female TBI+Ex rats treated with PRZ on day 49 post-TBI (**Fig 6A**). For assessment at 180 DPI, a cross over design was adopted. Female TBI+Ex rats that had been treated with PRZ on day 49 post-TBI were treated with the VEH at 180 DPI and the TBI+Ex rats treated with the VEH on day 49 after injury were treated with PRZ at 180 DPI. The DCN response was also blocked in all female TBI+Ex rats treated with PRZ on day 180 post-TBI (**Fig 6B**). VEH had no effect on the DCN response in female TBI+Ex rat at either timepoint (**Fig 6A** and **6B**). At 49 DPI a four-way mixed repeated measures ANOVA revealed significant main effects of drug condition (F(1,10)=1076.66, p=1.63 x 10^-11^, □2=0.99), and time post-capsaicin (F(7,70)=108.30, p=5.55 x 10^-35^, □2=0.92). There was also a significant interaction of time post-capsaicin by drug condition (F(7,70)=105.89, p=1.13 x 10^-34^, □2=0.91). At 180 DPI a four-way mixed repeated measures ANOVA revealed significant main effects of drug condition (F(1,10)=1459.41, p=3.60 x 10^-12^, □2=0.99), and time post-capsaicin (F(7,70)=137.94, p=2.31 x 10^-38^, □2=0.93). There was also a significant interaction of time post-capsaicin by drug condition (F(7,70)=134.92, p=4.74 x 10^-38^, □2=0.93). ⋇ = TBI+Ex+PRZ versus TBI+Ex+VEH (p < 0.0001) by Bonferroni’s posthocs. Open arrow - PGE-2; i.pl. [1 μg/50 μl]. Closed arrow - Capsaicin; i.pl. [10 μg/10 μl]. Data are presented as mean ± standard error of the mean. (n = 6 rats per cohort).

### The presence of a DCN response in male exercised-TBI rats is not dependent on intact 5-HT_7_ receptor signaling

For male TBI rats, we previously demonstrated that pharmacologic restoration of the DCN response is dependent on enhancing spinal serotonin levels to promote descending serotoninergic pain inhibition via the spinal 5-HT_7_ receptor^17,18^. Agents raising noradrenalin levels alone are ineffective^13,18^. To ascertain if this serotonergic mechanism is responsible for the restoration of the DCN response in exercised male rats (TBI+Ex), we treated them with the 5-HT_7_ receptor antagonist, SB-269970 shown effective in blocking the restoration of DCN after TBI by serotonin reuptake inhibitors in male rats^17^. Treatment began an hour after the conditioning stimulus (PGE-2), and SB-269970 had no effect on hindpaw sensitivity 15 minutes after its injection *prior* to the test stimulus (**Fig. 7: SB-26**). Surprisingly however, following the application of the test stimulus, SB-269970 failed to block the DCN response in the male TBI+Ex males at 49 DPI (**Fig. 7**). A four-way mixed repeated measures ANOVA revealed significant main effects of time post-capsaicin (p=1.17 x 10^-31^). No other main effects or interactions reached significance.

**FIGURE 7:**
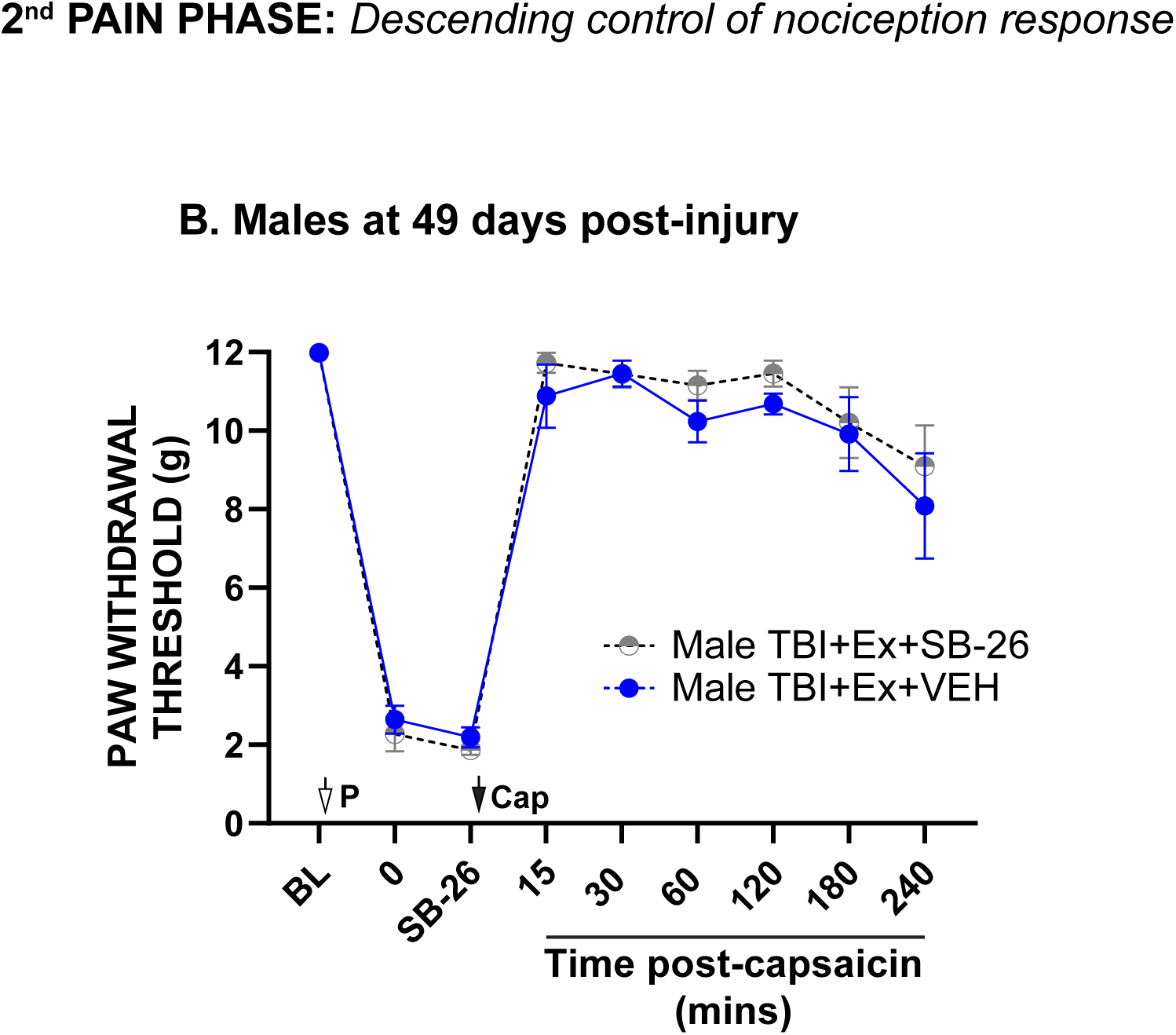
The effect of the 5-HT_7_ receptor antagonist, SB-267790 (SB-26), on the intact DCN response in male TBI+Ex rats 49 DPI (Fig 7). On the day of testing, half of the male TBI+Ex rats were randomly selected and given either an intrathecal injection of SB-26 (10 μg/10 μl, i.t.) or saline, the vehicle (VEH) after the test stimulus (PGE-2). The DCN response remained intact in male TBI+Ex rats treated with SB-26 at 49 DPI (Fig 7). VEH had no effect on the DCN response in male TBI+Ex rats. A four-way mixed repeated measures ANOVA revealed significant main effects of time post-capsaicin (F(7,70)=84.92, p=1.17 x 10^-31^, □2=0.90). *Post hoc* analysis reveals no significant difference in pain thresholds between TBI+Ex+SB-26 and TBI+Ex+VEH male rats. Data are presented as mean ± standard error of the mean. (n = 6 rats per cohort).

### The presence of a DCN response in male exercised-TBI rats is also dependent on intact α1 adrenoceptor signaling at days 49 and 180 post-injury

To establish the mechanism responsible for the presence of a DCN response in male TBI+Ex rats, we next treated them with prazosin using the same protocol as above for the female TBI+Ex rats. Systemic treatment with prazosin had no effect on level of hindpaw sensitivity induced by the conditioning stimulus (PGE-2) (**Fig. 8A: PRZ** and **8B: PRZ**). Following application of the test stimulus, the DCN response in the male TBI+Ex rats treated with prazosin was absent when tested both at 49 and 180 DPI (**Fig. 8A** and **8B**). Thus, similarly to the female TBI+Ex rats the mechanism responsible for the DCN response in the male TBI+Ex rats was α1AR dependent. At 49 DPI a four-way mixed repeated measures ANOVA revealed significant main effects of drug condition (p=4.83 x 10^-21^), and time post-capsaicin (p=5.69 x 10^-70^). There was also a significant interaction of time post-capsaicin by drug condition (p=1.00 x 10^-69^) and time post-capsaicin by sex (female, Figure 6 versus male, Figure 8) (p=0.0293). There was no significant main effect of sex or sex by drug condition interactions, all Ps greater than 0.05.

**FIGURE 8:**
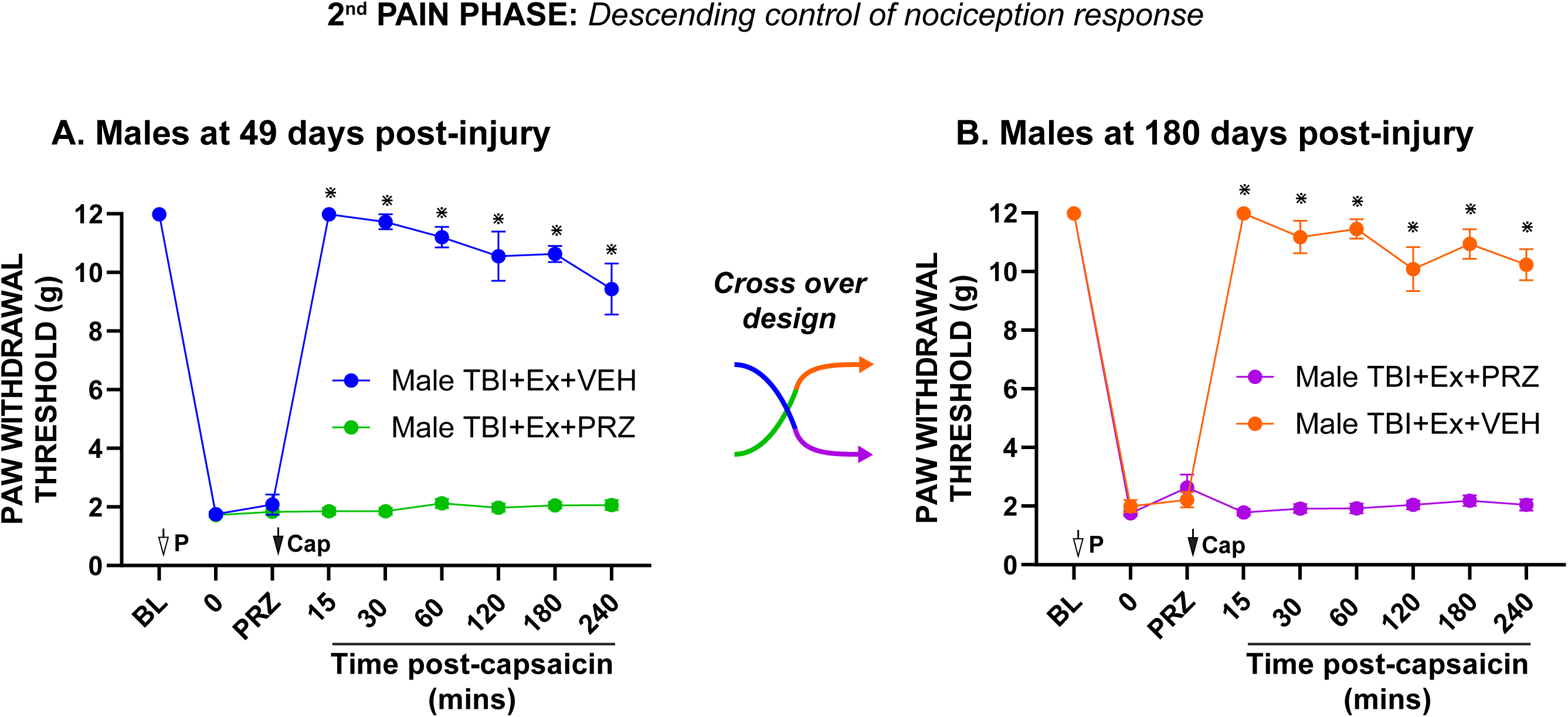
The effect of the α_1_-adrenoceptor (α_1_-AR) antagonist, prazosin (PRZ) on the intact DCN response in male TBI+Ex rats 49 (**Fig 8A**) and 180 DPI (**Fig 8B**). On day 49 of testing, half male TBI+Ex rats were randomly selected and given a systemic injection of PRZ or saline, the vehicle (VEH) (**Fig 8A** and **8B**) after the test stimulus (PGE-2). The DCN response was blocked in all male TBI+Ex rats treated with PRZ on day 49 post-TBI (**Fig 8A**). For assessment at 180 DPI, a cross over design was adopted. Male TBI+Ex rats that had been treated with PRZ on day 49 post-TBI were treated with the VEH at 180 DPI and the TBI+Ex rats treated with the VEH on day 49 after injury were treated with PRZ at 180 DPI. The DCN response was also blocked in all male TBI+Ex rats treated with PRZ on day 180 post-TBI (**Fig 8B**). VEH had no effect on the DCN response in male TBI+Ex rat at either timepoint (**Fig 8A** and **8B**). A four-way mixed repeated measures ANOVA comparing *both* male and female TBI+Ex PRZ and VEH treated rats at 49 DPI revealed significant main effects of drug condition (F(1,20)=1789.08, p=4.83 x 10^-21^, □2=0.99), and time post-capsaicin (F(7,140)=202.66, p=5.69 x 10^-70^, □2=0.91). There was also a significant interaction of time post-capsaicin by drug condition (F(7,140)=193.69, p=1.00 x 10^-70^, □2=0.91) and time post-capsaicin by sex (F(7,140)=2.31, p=0.0293, □2=0.10). There were no main effects of sex or sex by drug interactions found. At 180 DPI a four-way mixed repeated measures ANOVA comparing *both* male and female TBI+Ex PRZ and VEH treated rats revealed significant main effects of drug condition (F(1,20)=2546.36, p=1.46 x 10^-22^, □2=0.99), and time post-capsaicin (F(7,140)=185.55, p=1.50 x 10^-67^, □2=0.90). There was also a significant interaction of time post-capsaicin by drug condition (F(7,140)=193.27, p=1.15 x 10^-68^, □2=0.91) and time post-capsaicin by drug condition (F(7,140)=2.31, p=0.0295, □2=0.10). There were no main effects of sex or sex by drug interactions found. ⋇ = TBI+Ex+PRZ versus TBI+Ex+VEH (p < 0.0001) by Bonferroni’s posthocs. Open arrow - PGE-2; i.pl. [1 μg/50 μl]. Closed arrow - Capsaicin; i.pl. [10 μg/10 μl]. Data are presented as mean ± standard error of the mean. (n = 6 rats per cohort).

At 180 DPI a four-way mixed repeated measures ANOVA revealed significant main effects of drug condition (p=1.46 x 10^-22^), and time post-capsaicin (p=1.50 x 10^-67^). There was also a significant interaction of time post-capsaicin by drug condition (p=1.15 x 10^-68^) and time post-capsaicin by sex (p=0.0295). There was no significant main effect of sex or sex by drug condition interactions, all Ps greater than 0.05. *Post hoc* analysis reveals that the male TBI+Ex+prazosin groups had significantly lower pain thresholds than the vehicle-treated TBI+Ex male rats (*p < 0.001).

## Neuropathology

### Exercise protects from TBI-induced axonal loss at 7 days post-injury

At 7 days post-injury (DPI), the extent of axonal loss after injury was assessed within the medial corpus callosum of both male and female TBI+Ex, and TBI+Sed rats compared to shams of the same sex (**Fig. 9A**: red box). Intact and healthy axons were identified using the phosphorylated neurofilament marker, SMI 31R (**Fig. 9B**). Post hoc analysis indicated that exercise significantly decreased the extent of axonal loss in both male and females compared to sedentary rats at 7 DPI. In fact, there was no significant difference between female TBI+Ex compared to female sham rats. However, axonal loss in the male TBI+Ex rats was significantly greater when compared to female TBI+Ex rats at 7 DPI (**Fig. 9C**). Furthermore, there was significantly more axonal loss in the male TBI+Sed rats when compared to female TBI+Sed rats. Therefore, exercise protected against axonal loss in both male and female TBI rats. However, the overall extent of axonal loss was significantly greater for both exercise and sedentary TBI male rats compared to the female counterparts (**Fig. 9C**). A two-way mixed repeated measures ANOVA revealed significant main effects of exercise (p=1.14 x 10^-10^), and sex (p=1.47 x 10^-9^). There was also a significant interaction of group by sex (p=1.14 x 10^-4^). *Post hoc* analysis reveals that the male and female TBI groups have significantly less SMI 31R positive axons than the sham groups of the same sex (• p < 0.05). Male and female TBI+Ex groups have significantly more SMI 31R positive axons compared to male and female TBI+Sed groups of the same sex (# p < 0.05). Female TBI groups have significantly more SMI 31R positive axons than the male TBI groups (****p < 0.001).

**FIGURE 9:**
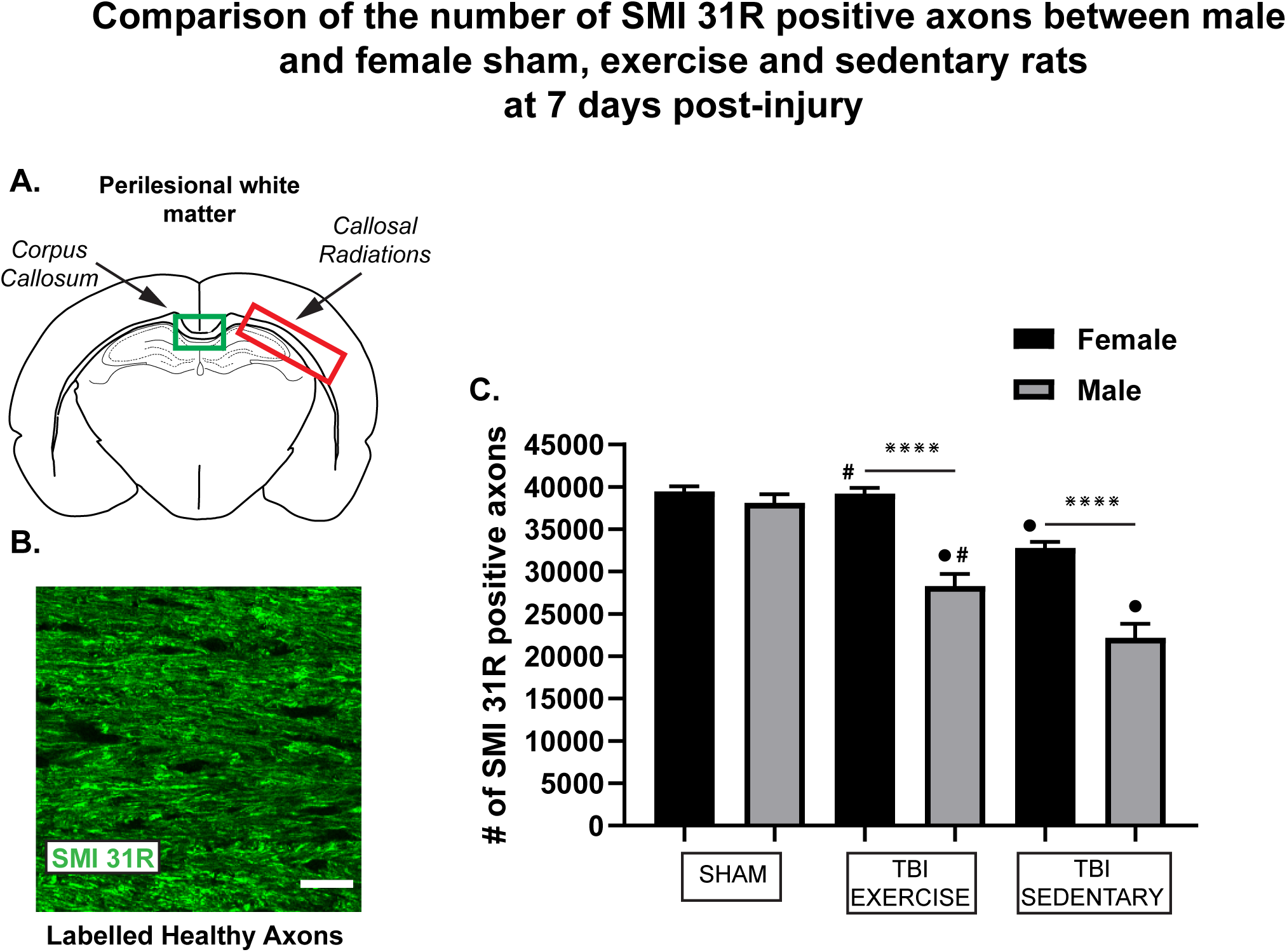
A comparison of the extent of axonal loss between male and female sham, TBI+Ex and TBI+Sed rat groups at 7 days post-injury (DPI). The number of intact and healthy axons were quantified at the medial corpus callosum (MCC) (**Fig 9A**: green box) using the phosphorylated neurofilament marker, SMI 31R (**Fig 9B**). Location of the callosal radiations is also indicated by the red box in **Fig 9A**. There was no significant difference in the number of SMI31R positive axons between female sham and female TBI+Ex rats (**Fig 9C**). In contrast, there was significant axonal loss in the male TBI+Ex rats compared to both male shams and female TBI+Ex rats (**Fig 9C**). Both male and female TBI+Sed rats had significantly less axons in the MCC when compared to male and female TBI+Ex rats respectively (**Fig 9C**). Furthermore, the extent of axonal loss in the male TBI+Sed rats was significantly greater when compared to the female TBI+Sed rats (**Fig 9C**). Two-way mixed repeated measures analysis of variance revealed a significant main effect of exercise (F(2,30)= 54.02, P= 1.14x 10^-10^, □2= 0.78) and sex (F(1,30)= 73.45, P= 1.47 x 10^-9^, □2= 0.71) and a group by sex interaction (F(2,30)= 12.47, P= 1.14 x10^-4^, □2= 0.45). #: Exercise TBI group vs. Sedentary TBI group of same sex, •: TBI group vs. Sham group of same sex) □=Female TBI vs. male TBI (p< 0.001), ⋇⋇⋇⋇: female vs. male (p< 0.0001), by Bonferroni’s posthocs. Data are presented as mean ± S.E.M. (n = 6, all groups).

### Effect of exercise on the microglial/macrophage response to TBI at 7 days post-injury

The effect of exercise on the microglial/macrophage population was assessed using IBA-1 immunohistochemistry of several regions of the brain: rostral ventral medulla (RVM), ventrolateral periaqueductal grey (vlPAG), locus coeruleus (LC), thalamus (TH), dentate gyrus (DG), perilesional grey (CTX) and white matter (CR) (**Fig. 10** and **Table 1**). Post hoc analysis indicated that exercise significantly increased IBA-1 expression on the ipsilateral side of the CTX, CR, TH, DG and the LC in female rats when compared to female TBI+Sed rats (**Fig. 10A to 10E**). In contrast, an exercise induced *increase* in IBA-1 expression was only present in the ipsilateral TH and DG of male TBI rats compared to male TBI+Sed rats (**Fig. 10C and 10D**). In the LC of male TBI+Ex rats there was a significant decrease bilaterally in IBA-1 expression compared to male TBI+Sed (**Fig. 10E**). No further exercised-induced changes to the expression of IBA-1 were visible in either male or female TBI rats (**Fig. 10**). Significant main effects and interactions from a three-way mixed repeated measures ANOVA are displayed in **Table 1.** *Post hoc* analysis reveals significantly greater IBA-1 expression compared to the sham group of the same sex (• p < 0.05). Significantly different % of IBA-1 expression on the ipsilateral side of the region of interest compared to the contralateral side (# p < 0.05). Significantly different % of IBA-1 expression in the TBI+Ex group compared to the TBI+Sed group of the same sex (◊ p < 0.05). Significantly different % of IBA-1 expression between male TBI group and female TBI group (* p < 0.05, **p < 0.01, ***p < 0.001, ****p < 0.0001).

**FIGURE 10:**
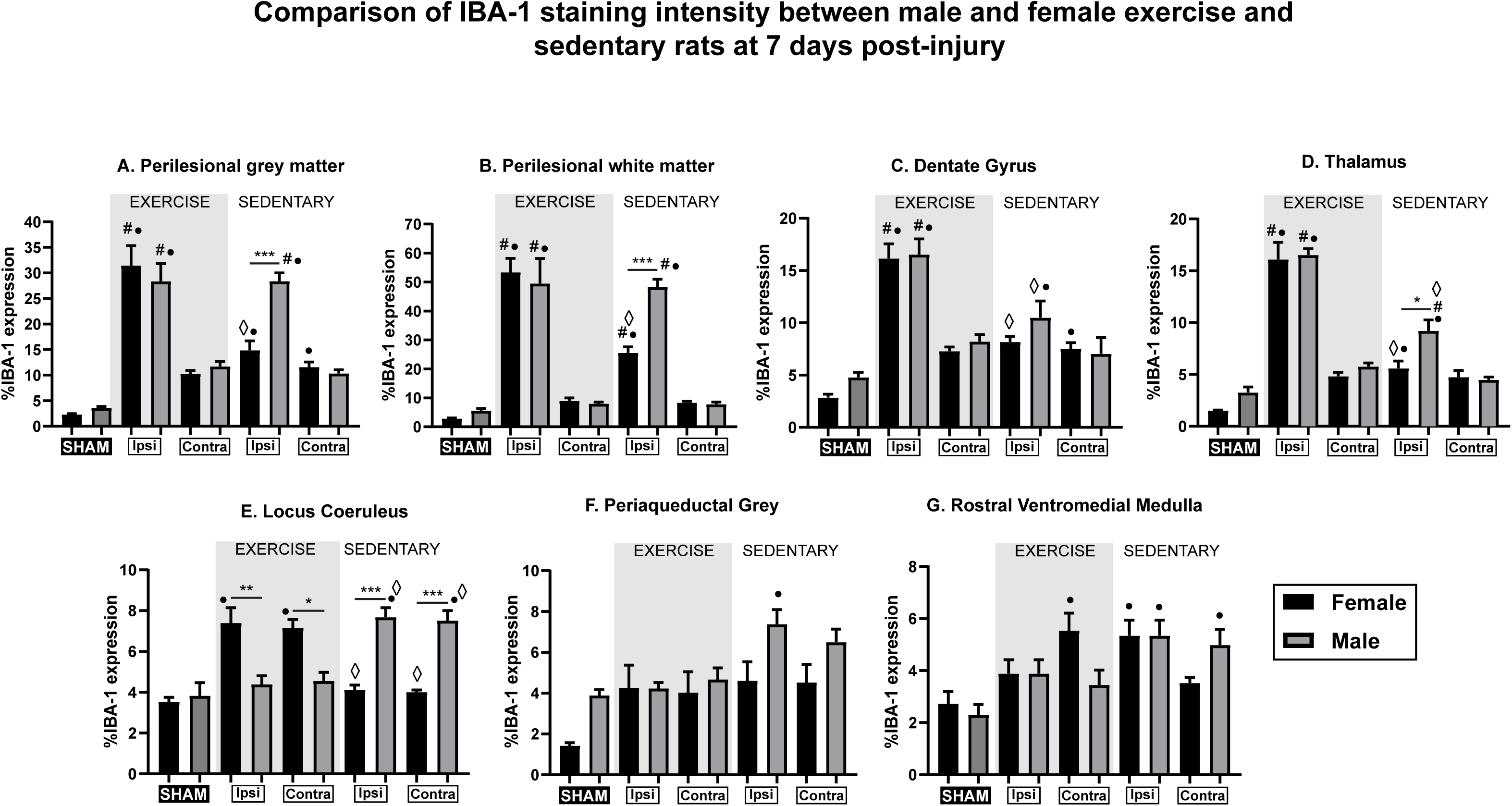
The effect of exercise on the expression of IBA-1 (for microglia and macrophages) in several regions of the brain including the perilesional grey matter (**Fig 10A**: cortex) and white matter (**Fig 10B**: callosal radiations), dentate gyrus (**Fig 10C**), thalamus (**Fig 10D**) and the pain centers the locus coeruleus (**Fig 10E**), periaqueductal grey (**Fig 10F**) and rostral ventromedial medulla (**Fig 10G**). Groups compared include male and female, sham, TBI+Exercise and TBI+Sedentary rats. ◊: Exercise vs sedentary of same sex (p< 0.05), #: Ipsilateral side vs Contralateral side of same sex (p< 0.05), •: TBI group vs. Sham group of same sex (p< 0.05), * = (p< 0.05), ** = (p< 0.01), *** = (p< 0.001) and **** = (p< 0.0001) by Bonferroni’s posthocs. Values are displayed as mean ± standard error of the mean. n = 6 rats/cohort.

**Table 1:**
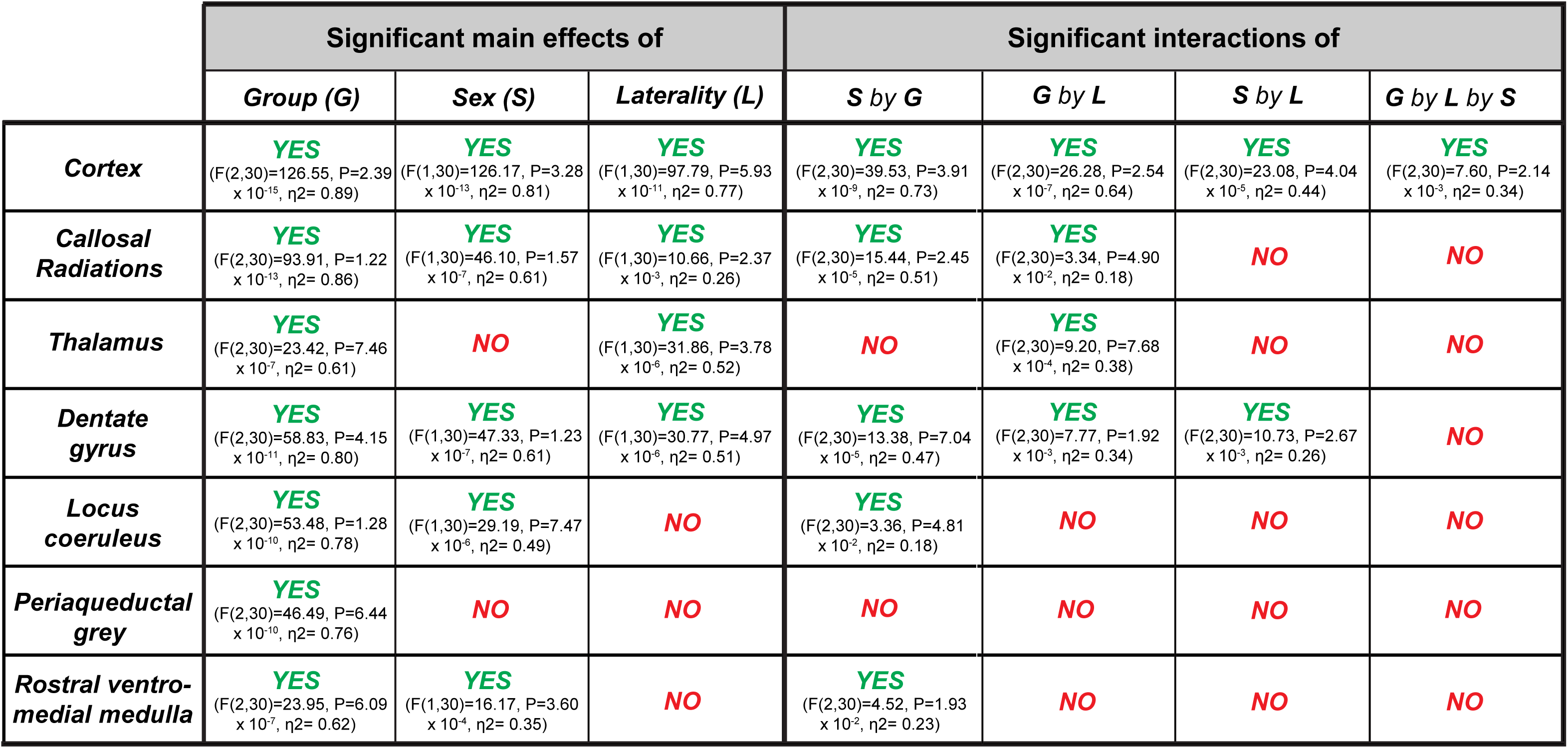
This table displays the results from the 4-way mixed repeated measures analysis of the % intensity of GFAP staining of the brain for astrocytes at 7 days post-injury. Results were split into significant main effects of and significant interactions between group, sex, and laterality (ipsilateral side versus contralateral side). Data are presented are the precise F values, degrees of freedom, P values, effect sizes, and observed power for all significant effects. (n = 6, all groups).

### Effect of exercise on the astrocyte response to TBI at 7 days post-injury

The effect of exercise on the astrocyte population was assessed using GFAP immunohistochemistry of several regions of the brain: rostral ventral medulla (RVM), ventrolateral periaqueductal grey (vlPAG), locus coeruleus (LC), thalamus (TH), dentate gyrus (DG), perilesional grey (CTX) and white matter (CR) (**Fig. 11** and **Table 2**). Post hoc analysis indicated that exercise decreased GFAP expression in the ipsilateral DG compared to female TBI+Sed rats (**Fig. 11C**). No additional effects of exercise on GFAP expression in female TBI rats were present (**Fig. 11**). In male TBI+Ex rats exercise decreased the expression of GFAP in the ipsilateral CTX (**Fig. 11A**). No further exercise-induced changes on GFAP expression were present in male TBI in the other regions analyzed. When the effect of exercise on GFAP expression was compared between male and female TBI rats it revealed that there was a significantly greater GFAP expression in male TBI+Ex in the ipsilateral CTX, CR, TH, LC and RVM compared to female TBI+Ex rats (**Fig. 11A-B, D, E and G**). Significant main effects and interactions from a three-way mixed repeated measures ANOVA are displayed in **Table 2.** *Post hoc* analysis reveals significantly greater GFAP expression compared to the sham group of the same sex (• p < 0.05). Significantly different % of GFAP expression on the ipsilateral side of the region of interest compared to the contralateral side (# p < 0.05). Significantly different % of GFAP expression in the TBI+Ex group compared to the TBI+Sed group of the same sex (◊ p < 0.05). Significantly different % of GFAP expression between male TBI group and female TBI group (* p < 0.05, **p < 0.01, ***p < 0.001, ****p < 0.0001).

**FIGURE 11:**
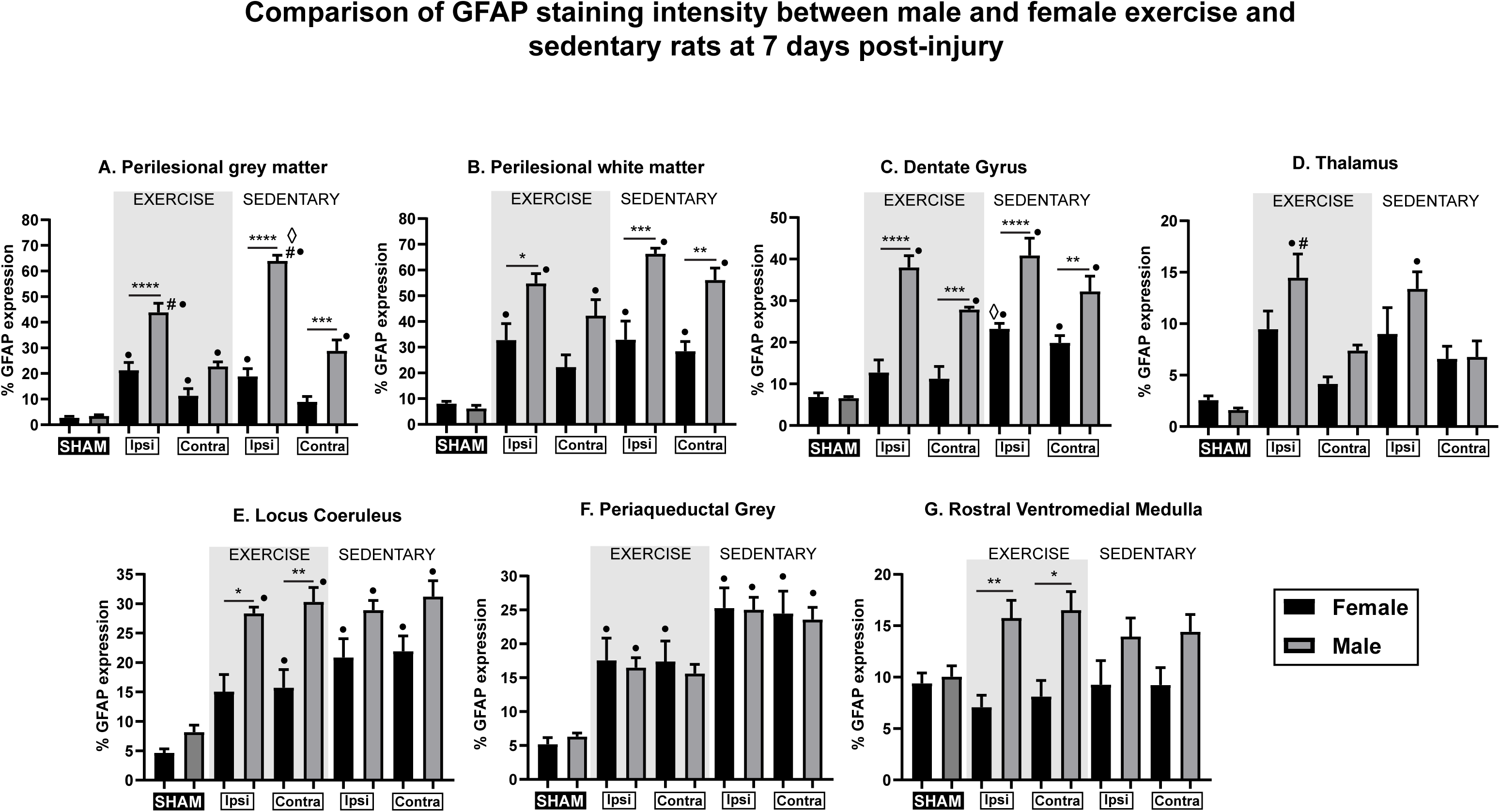
The effect of exercise on the expression of GFAP (for astrocytes) in several regions of the brain including the perilesional grey matter (**Fig 11A**: cortex) and white matter (**Fig 11B**: callosal radiations), dentate gyrus (**Fig 11C**), thalamus (**Fig 11D**) and the pain centers the locus coeruleus (**Fig 11E**), periaqueductal grey (**Fig 11F**) and rostral ventromedial medulla (**Fig 11G**). Groups compared include male and female, sham, TBI+Exercise and TBI+Sedentary rats. ◊: Exercise vs sedentary of same sex (p< 0.05), #: Ipsilateral side vs Contralateral side of same sex (p< 0.05), •: TBI group vs. Sham group ofsame sex (p< 0.05), * = (p< 0.05), ** = (p< 0.01), *** = (p< 0.001) and **** = (p< 0.0001) by Bonferroni’s posthocs. Values are displayed as mean ± standard error of the mean. n = 6 rats/cohort.

**Table 2:**
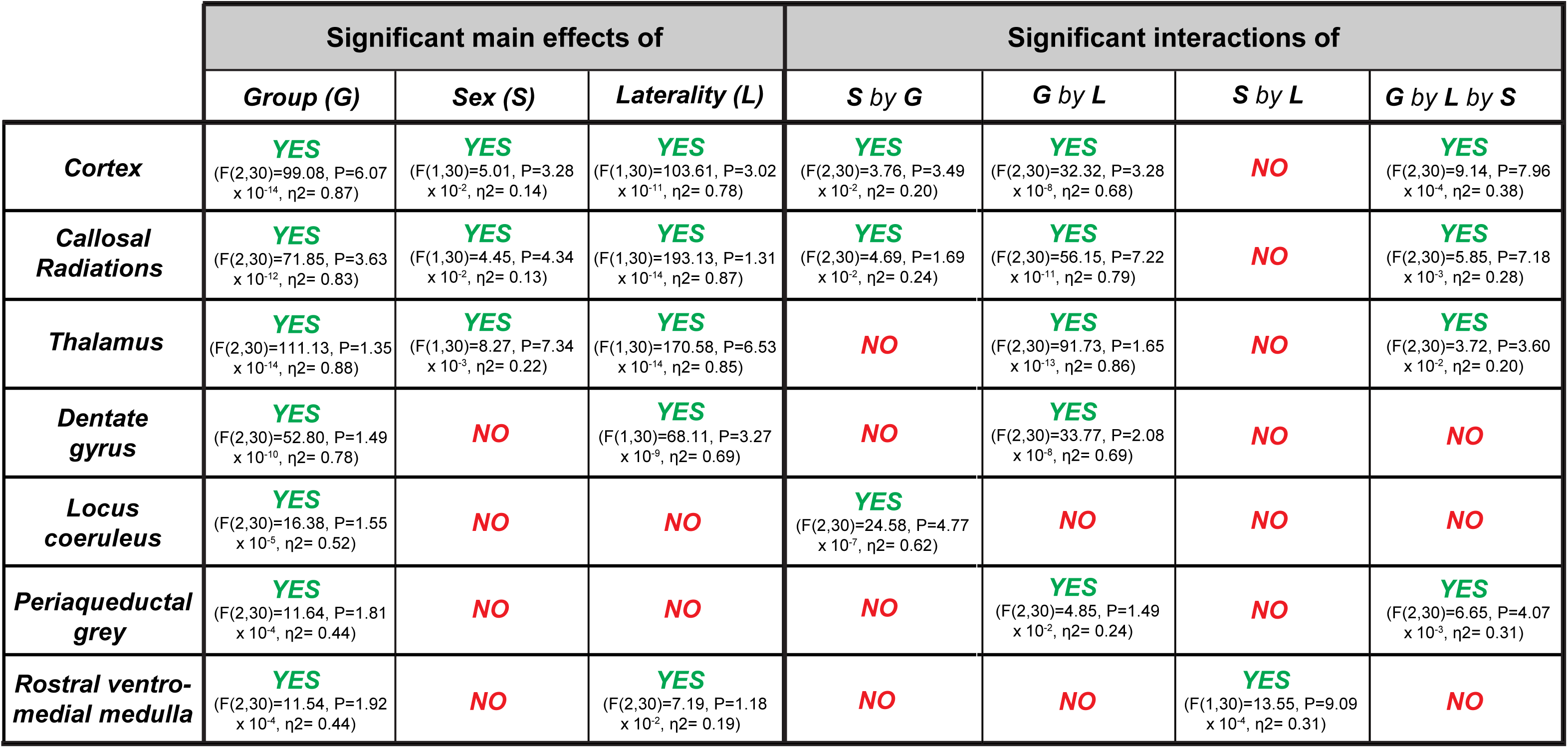
This table displays the results from the 4-way mixed repeated measures analysis of the % intensity of IBA-1 staining of the brain for microglia/macrophages at 7 days post-injury. Results were split into significant main effects of and significant interactions between group, sex, and laterality (ipsilateral side versus contralateral side). Data are presented are the precise F values, degrees of freedom, P values, effect sizes, and observed power for all significant effects. (n = 6, all groups).

## Discussion

Chronic pain is one of the most significant barriers to recovery after TBI which can thwart rehabilitation efforts, cause emotional distress and worsen cognitive deficits. Deficiencies in endogenous pain modulation are suggested to be responsible for the development of chronic pain in a wide variety of conditions such as osteoarthritis, fibromyalgia, opioid-induced hyperalgesia and post-traumatic headache (PTH). We and others have shown that the reduction in descending pain inhibition and the enhancement of descending facilitation of pain is responsible for the manifestation of chronic pain after TBI^5^. Occurring at significantly higher rates in mild TBI survivors compared to moderate to severely injured TBI patients, the most prevalent form of chronic pain is PTH followed closely by back, joint, and other forms of musculoskeletal pain^32^.

Aerobic exercise is characterized by continuous and rhythmic movements that result in both an increase in heart rate and oxygen intake such as swimming, walking, and cycling. The positive correlation between aerobic exercise and TBI recovery in animal studies (enhancing memory, neuroplasticity, and lowering anxiety) is well documented whilst the role of exercise and TBI in human patients is still in its early stages^33^. Those studies that do exist largely suggest that exercise following a TBI improves the patients “quality of life”, an outcome measure based upon the patient’s functional and mental health status, and psychosocial function^34,35^. Large population studies show that people who are physically active report a lower incidence of chronic pain^36,37^. TBI patients have reported that physical activity is linked to lower pain one month after mild TBI, effects that may be linked to more efficient endogenous pain modulation^38^.

In this study we investigate the effect of voluntary wheel running exercise on TBI-induced pain in both male and female Sprague-Dawley rats after a lateral fluid percussion injury. Our first observation into the effect of exercise on pain outcome measures after TBI revealed that it can significantly reduce the duration of the acute stage of pain from 4-5 weeks to 2 to 3 weeks in female and male TBI rats respectively. Exercise-induced restoration of the acute stage in both male and female TBI rats could be reversed using the α_1_-AR antagonist, prazosin (**Fig. 3**). So, exercise did not prevent the switch from the normally operative α_2_-AR system to α_1_-AR mediated descending inhibition. Assessment at 49 and 180 DPI revealed that exercise prevented the loss of the DCN response in both male and female rats when compared to sedentary TBI rats. The exercised induced “protection” of the DCN response in female TBI rats after injury could be reversed using prazosin. Surprisingly, the intact DCN response in the male TBI+Ex rats was not blocked by the 5-HT_7_ receptor antagonist, SB-269970 as expected based on our previous observations of this receptor being involved in restoration of DCN in male but not female TBI rats^13,17^. In fact, following exercise the DCN response in male TBI rats was blocked using prazosin similar to the females.

We have previously shown that acutely after TBI (7 DPI) descending NA inhibition of pain in both male and female rats switch from an α_2_-AR to an α_1_-AR mediated pathway^13^. While females remain dependent on NA signaling via the α_1_-AR up to 180 DPI, males, at some point prior to 49 DPI, switch to a purely serotonergic driven pain inhibitory pathway via 5-HT_7_^13,18^. When and why, this occurs is subject of future exploration. However, what is clear is that exercise blocked this transition in the male TBI+Ex rats.

One explanation for this could be exercise-induced protection of the LC. TBI has been consistently found to cause the loss of neurons (45%) in and projections (65%) from the LC. This results in the loss of both central NA levels and NA turnover after injury^39^ ^40^. Axons of LC neurons are unmyelinated which makes them very susceptible to mechanical injury after TBI^41^. Together, this suggests that the LC is a highly vulnerable structure to damage after TBI. Interestingly, the LC in female rats has been shown to be larger and contain significantly more NA containing neurons when compared to male rats ^42–44^. Furthermore, dendrites of the LC neurons of female rats were more complex and extended further covering a significantly greater anatomical area than those of the males^45^. It is possible that due its smaller size and that it contains fewer NA positive neurons, that the male LC is more susceptible to damage after TBI. Damage to the LC may be extreme enough in male TBI rats that the descending NA inhibitory pain pathway simply becomes nonfunctional. This then leaves only the descending serotonergic inhibitory pathway remaining in male TBI rats. Exercise has been shown to activate the LC, increase the release of NA and provide galanin to promote hyperpolarization of the LC neurons^46–49^. During times of injury, hyperpolarization is protective and would reduce the susceptibility of LC neurons to stress and neurodegeneration. Therefore, exercise may have protected the LC, in male TBI rats, from TBI induced neurodegeneration sufficiently enough to protect the integrity of the descending NA inhibitory pain pathway.

Exercise also significantly decreased the extent of axonal loss in the corpus callosum (CC), the white matter tract closest to the lesion site in both male and female TBI rats. However, when male and female TBI rats were compared, the extent of callosal axon loss was significantly higher in male compared to female rats, regardless of exercise.

Analysis of the microglial and astrocyte response revealed significantly higher expression of both IBA-1 and GFAP in the CTX, CR and TH of male TBI+Sed compared to female TBI+Sed rats. In contrast, there was no significant difference in IBA-1 expression in these areas between male and female TBI+Ex rats. Unexpectedly however, the expression of IBA-1 was significantly lower in the TBI+Sed rats compared to the TBI+Ex rats for both sexes. GFAP expression was significantly higher in the CTX, CR and DG of male TBI+Ex compared to female TBI+Ex rats. However, exercise appeared to have little if no effect in any regions analyzed apart from the CTX and DG in male and female TBI rats respectively.

Neuroinflammation is considered to be a proverbial “double edge sword” meaning that the presence of neuroinflammation following injury is critical for repair processes after TBI^50^. However, if left unresolved, can lead to the overproduction and release of multiple mediators including pro-inflammatory cytokines (IL-1β, TNFα),^51^ that can increase the extent of brain damage ^52^. Exercise can have a number of effects on neuroinflammation after brain injury that may not necessarily be reflected in a change in expression GFAP and IBA-1. Exercise has been shown to reduce the expression of proinflammatory cytokines, modulate cell activity to promote a pro-repair phenotype, increase the release of anti-inflammatory cytokines (IL-4, −10 and −13) and enhance cell survival and plasticity via the release of neurotrophic factors such as NGF and BDNF^53–57^. Therefore, replication of this study with a focus on expression of inflammatory cytokines, chemokines and neurotrophic factors would be a logical next step.

For this study, exercise began on day 3 and continued until day 49 after TBI. Surprisingly, despite its cessation at 49 DPI, the effects of exercise on the DCN response were still present ∼4 months later on day 180. Therefore, the effect of exercise on the DCN response was persistent if not permanent. A suitable follow-up study would be to terminate exercise at several timepoints prior to 49 DPI to observe if the minimum number of days of exercise required to preserve the DCN response can be reduced.

There are limitations to this study. For example, the estrus cycle of the female rats used in this study was not monitored prior to injury. The estrus cycle, is characterized by 4 distinct phases (proestrus, estrus, metestrus, and diestrus)^58^ during which female sex hormones estrogen and progesterone circulate and due to their lipophilic nature, can cross the blood brain barrier and alter neuroactivity of the brain. TBI experiments that vary when the female rats obtain an injury relative to the estrus cycle have shown disparities in neurocognitive and psychological outcomes^59–62^. Replicating this study and controlling for the phase of the estrus cycle at the time of injury will be necessary to confirm our current findings.

In conclusion, our results demonstrate that voluntary running exercise in SD rats after LFP injury significantly reduces the duration of the acute pain stage from ∼5 weeks to 2 to 3 weeks in female and male TBI rats respectively. Restoration of normal hindlimb sensitivity in both male and female TBI+Ex rats could be reversed using the α_1_-AR antagonist, prazosin. Assessment of the endogenous nociceptive control systems during the chronic stage of pain revealed that exercise had prevented the TBI-induced loss of the DCN response in both male and female TBI+Ex rats. The DCN response in TBI+Ex rats, regardless of sex, could be blocked using prazosin. This is consistent to what we have shown previously in female TBI rats however, in male TBI rats the spinal noradrenergic analgesic systems become non-functional. Restoration of the DCN response is still possible in male TBI rats due to the transition to a serotonergic, specifically 5-HT_7_ dependent mechanism. Therefore, exercise appears to have blocked the transition from descending NA to 5-HT inhibitory pain signaling in male TBI+Ex rats. A key finding in the assessment of the neuroinflammatory response between male and female TBI rats revealed that both IBA-1 and GFAP expression was significantly higher in male TBI rats compared female TBI rats, regardless of exercise. Whereas the difference in the expression of IBA-1 and GFAP in TBI+Ex compared to TBI+Sed rats was minimal for both males and females. In contrast, exercise significantly decreased the extent of axonal loss in the corpus callosum of both male and female TBI+Ex compared to TBI+Sed rats. Nevertheless, it was clear that there was significantly more axonal loss in male TBI rats compared to female TBI rats regardless of exercise.

### Transparency, Rigor, and Reproducibility Summary

All studies were approved by the Veterans Affairs Palo Alto Health Care System Institutional Animal Care and Use Committee (Palo Alto, CA, USA) and followed the animal guidelines of the National Institutes of Health Guide for the Care and Use of Laboratory animals (NIH Publications 8th edition, 2011). A total of 106 male and 124 female Sprague Dawley rats were randomly divided into different groups. An independent statistician (A.R.F.) tested statistical assumptions, performed missing values analysis, descriptive statistical tests, followed by inferential tests. Data met normality, homogeneity of variance, and independence assumptions sufficiently for application of general linear models (GLM). Group comparisons were therefore performed according to *a priori* balanced experimental designs as three-way, four-way, or five-way mixed ANOVAs, modeling between- and within-subject factors as appropriate using GLM. Significant effects were followed by interaction plots and Bonferroni’s post hocs. All statistical analyses were performed using IBM SPSS software (v.31, IBM Corp.) with base, regression, missing values, and advanced statistics modules. Statistical code is freely available upon request. Graphs were created using Prism 10.1.0 (GraphPad Software). Data are presented as mean values ± standard error of the mean (SEM). All experimental sample sizes (n) were selected by *a priori* power calculations based on historical data from our laboratory. The figure legends report precise F values, degrees of freedom, p values, effect sizes, and observed power for all significant effects as well as the n’s completed for each group. Experimenters were blinded to the identity of treatments and experimental conditions. All studies were designed to minimize the number of rats required. Before testing, all rats were also transferred into the testing room to acclimate to the environment for 60 min to ensure consistent activity. Mechanical withdrawal thresholds were measured using a modification of the up-down method and von Frey filaments as described previously^24^. Our investigators were proficiently trained in advance to use the von Frey apparatus.

## Authors’ Contributions

Karen-Amanda Irvine: conceptualization, methodology, investigation, data curation, simple analysis, original draft, review, and editing. Adam R Ferguson: formal analysis, review, and editing. J. David Clark: conceptualization, methodology, original draft, review, and editing.

## Acknowledgments

This work was supported by grants from the U.S. Dept. of Defense (MR130295) and the Dept. of Veterans Affairs (RX001776) and (2 I01 RX001776-05). ARF supported by grant from Dept. of Veterans Affairs (RX002787), (I50BX005878), (I01BX005871) and NIH/NINDS (UH3NS106899; U24NS122732).

